# Progressive Post-Acute Gut Microbiome Disruption Following SARS-CoV-2 Infection in Mice

**DOI:** 10.64898/2026.09.16.751817

**Authors:** Ritu Mann-Nüttel, Natacha S. Ogando, Marie Armbruster, Shivani Mandal, Dusadee Ospondpant, Mohamed Elaish, Tom C. Hobman, Christopher Power, Paul Forsythe

**Author notes:** **Corresponding author and email address:** Paul Forsythe. **Repositories:** Data is publicly available in the NCBI BioProject database (PRJNA1236398).

## Abstract

2.

SARS-CoV-2 infection is increasingly recognized to produce long-lasting physiological disturbances that extend beyond the acute phase. The gut microbiome has emerged as a potential contributor to post-infection outcomes, yet the longitudinal progression of microbial disruption following infection remains poorly defined. Here, we used a mouse model of SARS-CoV-2 infection to characterize gut microbiome dynamics at 7 (late acute phase), 14 (early recovery), and 21 (post-COVID condition, PCC) days post-infection (dpi). Viral RNA was detected in the lungs through 21 dpi. Longitudinal microbiome profiling revealed a two-phase restructuring marked by early shifts in Bacillota and Bacteroidota abundance, followed by pronounced post-acute dysbiosis characterized by reduced diversity, expansion of Bacillota, and near-complete loss of Bacteroidota. Functional pathway predictions showed progressive metabolic remodeling, including disruptions in nucleotide turnover, fermentation, and vitamin-related pathways. Together, these findings demonstrate that SARS-CoV-2 infection drives progressive and sustained alterations in gut microbial composition and function after the transition from acute infection through recovery into PCC.

**Data summary:** The authors confirm all supporting data, code and protocols have been provided within the article or through supplementary data files. Any other information can be acquired by contacting the first author via email.

## 4. Introduction

Coronavirus disease of 2019 (COVID-19), caused by severe acute respiratory virus 2 (SARS-CoV-2), has had a profound impact on global health. While much focus has been placed on prevention and the acute phase of infection, a growing body of evidence highlights long-term consequences, collectively termed post-COVID-19 condition (PCC). These lingering effects include fatigue, neurological symptoms, and immune dysregulation^1^, raising concerns about underlying biological mechanisms that persist beyond viral clearance. One emerging factor in this context is the alteration of gut microbiome, which plays a crucial role in immune modulation, metabolic function and host resilience^2^. Short-chain fatty acids (SCFAs), key microbial metabolites produced by commensal anaerobes, are particularly important for maintaining epithelial integrity, regulating inflammation, and supporting neuroimmune communication^3^.

SARS-CoV-2 infection has been associated with perturbations in the gut microbiota, including the depletion of beneficial commensal bacteria such as *Bifidobacterium*, *Lactobacillus*, and *Faecalibacterium prausnitzii*, alongside expansion of potentially opportunistic pathogenic species such as *Streptococcus pneumoniae* and *Haemophilus influenzae*^4,5^. These microbial shifts can influence systemic inflammation, intestinal permeability, and immune function, potentially contributing to some of the prolonged symptoms observed in PCC patients. A persistent dysbiosis of the gut microbiome may perpetuate symptoms like fatigue^6^ that are commonly observed in PCC. Microbiome dysbiosis may also promote chronic and systemic inflammation and immune dysfunction^7^. Moreover, there is evidence that communication between the gut microbiota and the brain influences cognitive function^8^, and an imbalance in microbiome composition may contribute to neurological symptoms like memory loss and concentration difficulties^9^. Given the central role of SCFA-producing taxa in these processes, disruptions to SCFA-generating microbial communities may represent a key mechanistic link between SARS-CoV-2 infection and PCC-related symptoms. Microbiome-targeted interventions including probiotics, prebiotics, and fecal microbiota transplantation are being explored as strategies to restore microbial balance in PCC patients^10,11^. However, mechanistic insight into SARS-CoV-2–induced dysbiosis remains limited, particularly regarding the temporal progression of microbial and metabolic alterations during and following infection. Overall, there is a need to better understand how SARS-CoV-2 disrupts the gut ecosystem and how these disruptions evolve over time.

Animal models, particularly murine models, have been instrumental in elucidating host–microbiome interactions during viral infections. Mouse models of COVID-19 have provided valuable insights into viral pathogenesis, immune responses, and post-infectious sequelae^12,13^. In these models, the peak of viral replication occurs at 2–4 days post-infection (dpi), with 7 dpi marking the end of the acute phase. Recovery begins around 14 dpi and extends into a prolonged period up to 21 dpi, which corresponds to PCC^14,15^. Recent mouse model studies have revealed pronounced late-phase neuroimmune disturbances, including persistent viral RNA in multiple brain regions and a shift from depressive-to anxiety-like behaviors in the post-infection period ^15^. These delayed central changes suggest that post-acute pathology continues to evolve well beyond the initial infection and raise the possibility that peripheral systems—including the gut microbiome—may undergo similar progressive remodeling. This evidence provided a rationale to investigate how the intestinal microbiome changes over the same post-acute timeline.

Thus, in this study, we utilized a mouse model of SARS-CoV-2 infection^16,17^ to characterize gut microbiome dynamics across key stages: 7 (end of acute phase), 14 (early recovery) and 21 dpi (PCC). By integrating microbiome composition and functional pathway predictions, we aimed to investigate the trajectory of gut microbial disruption following SARS-CoV-2 infection and identify microbial or metabolic features that may serve as potential therapeutic targets for microbiome-based interventions aimed at mitigating COVID-19 and PCC sequelae.

## 5. Results

### 5.1 Virological Profile and Microbiome composition at 7 dpi

Viral RNA levels were high in the lung at 7 dpi (18,061,250 ± 10,814,396 copies/g) but then dropped by ∼250-fold at 14 dpi (71,375 ± 26,882 copies/g) and by nearly 1,000-fold at 21 dpi (18,378 ± 6,474 copies/g), remaining low thereafter. (**Fig. 1A**). With the virological profile established, we next examined the gut microbiome at the corresponding time points. Baseline analysis of gut microbiome composition at 7 dpi revealed no significant differences in alpha-diversity metrics, including Shannon index, Chao1 richness, and Pielou evenness, between infected and mock-infected mice (p > 0.05) (**Fig. 1B**). Despite this overall similarity in within-sample diversity, there were clear phylum-level compositional differences between mock and infected groups. Bacillota A was the dominant phylum in both groups but was higher (71.3%) in infected than in mock-treated mice (57.4%). Conversely, Bacteroidota was decreased in infected (16.9%) compared to control (29.5%) mice (**Fig. 1C**). At the genus level, *Hominiventricola* abundance was significantly lower in infected animals at 7 dpi (**Fig. 1D**). Overall, these results demonstrate the disruptive effect of SARS-CoV-2 infection in the gut microbiome.

**Fig. 1.**
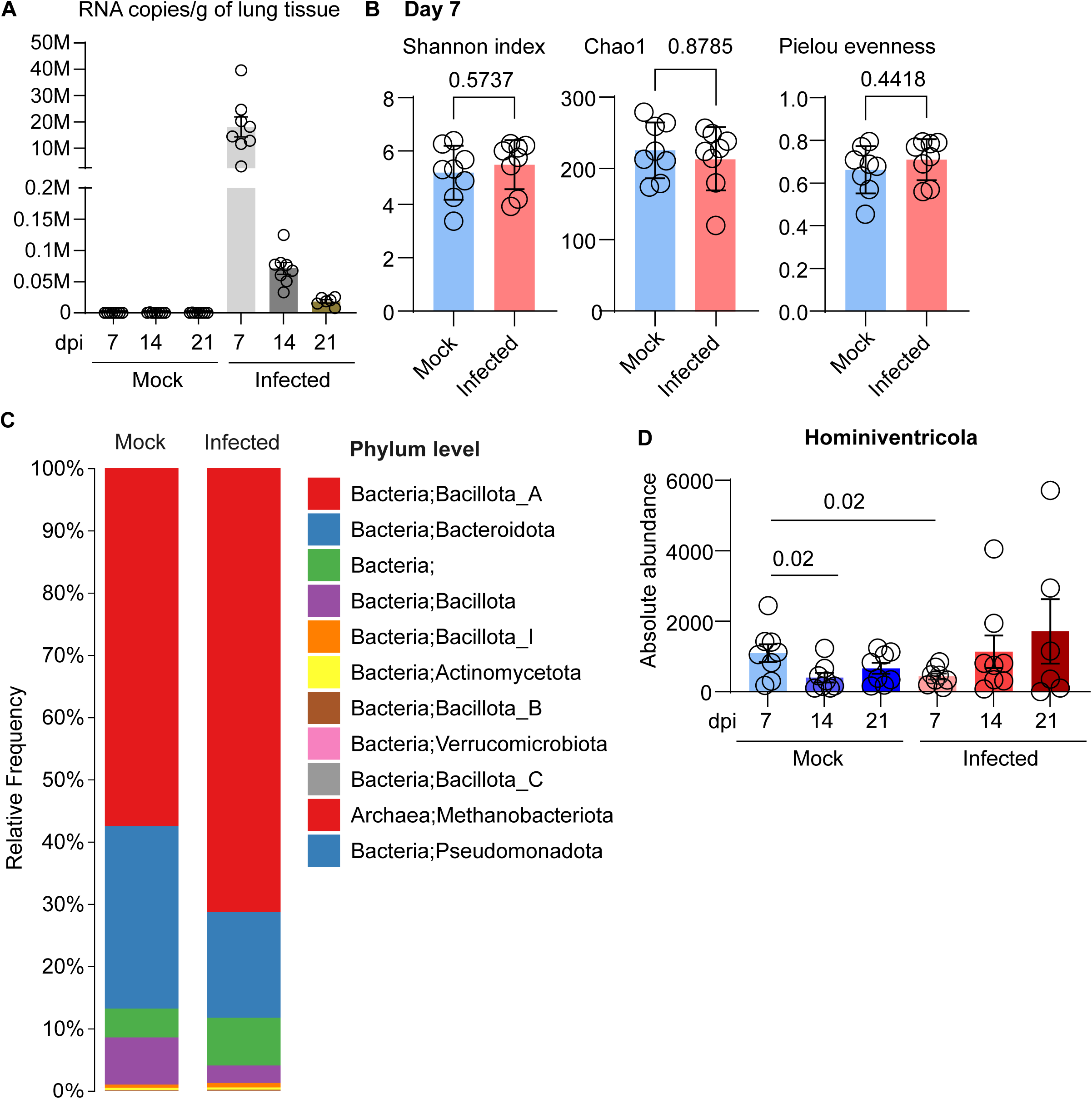
Baseline microbiome characterization at 7 dpi. **A** SARS-CoV-2 RNA copies/g of lung tissue at 7, 14 and 21 dpi for mock and infected. **B** Shannon index, Chao1 and Pielou Evenness (Alpha Diversity) for mock and infected at 7 dpi. **C** Mean relative frequencies of identified bacterial phyla across mock and infected samples at 7 dpi shown as percentage bar plots. **D** Absolute abundance level changes of Genus level bacteria between mock and infected at 7 dpi. Bar plots show individual values of biological replicates with error bars depicting SEM. Mann-Whitney (B) and Kruskal-Wallis (D) test performed for statistical analysis.

### 5.2 Longitudinal analysis reveals progressive changes in the microbiome on Genus level

Longitudinal analysis revealed a progressive increase in Bacillota A (Day 7: 71.32%; Day 14: 79.1%; Day 21: 91.6%) and a concurrent decline in Bacteroidota (Day 7: 16.9%; Day 14: 14.1%; Day 21: 3.7%) in infected mice over time (**Fig. 2A**). At the genus level, microbial shifts occurred exclusively in the infected group, with some bacterial taxa increasing (*Lachnospiraceae, Petralouisia, Acutalibacteraceae, MGBC164599*) while others decreased (*Alistipes, Hominilimicola, Butyribacter, Blautia A, Roseburia A, Tyzzerella*). Of note, all bacteria that showed significant changes in their absolute abundance belong to the phylum Bacillota A and the order of Clostridia except for *Alistipes*, that represents Bacteroidota. Looking at the time points we see two phases for microbiome changes: The first phase is the recovery time after the acute phase of infection, where *Alistipes, Hominillimicola, Lachnospiraceae* and *Petralouisia* and *MGBC164599* abundance changed comparing 7 and 14 dpi time points, while maintaining their levels up to 21 dpi. The phase after 14 dpi, which marks potentially the development of PCC phase, shows significant changes in absolute abundance of other taxa (*Butyribacter*, *Blautia A*, *Roseburia B*, *Tyzzerella* and *Acutalibacteraceae*) comparatively with 21 dpi (**Fig. 2B, C**). Importantly, these patterns were not observed in mock-infected mice, suggesting that SARS-CoV-2 infection contributes to the observed alterations in the microbiome composition.

**Fig. 2.**
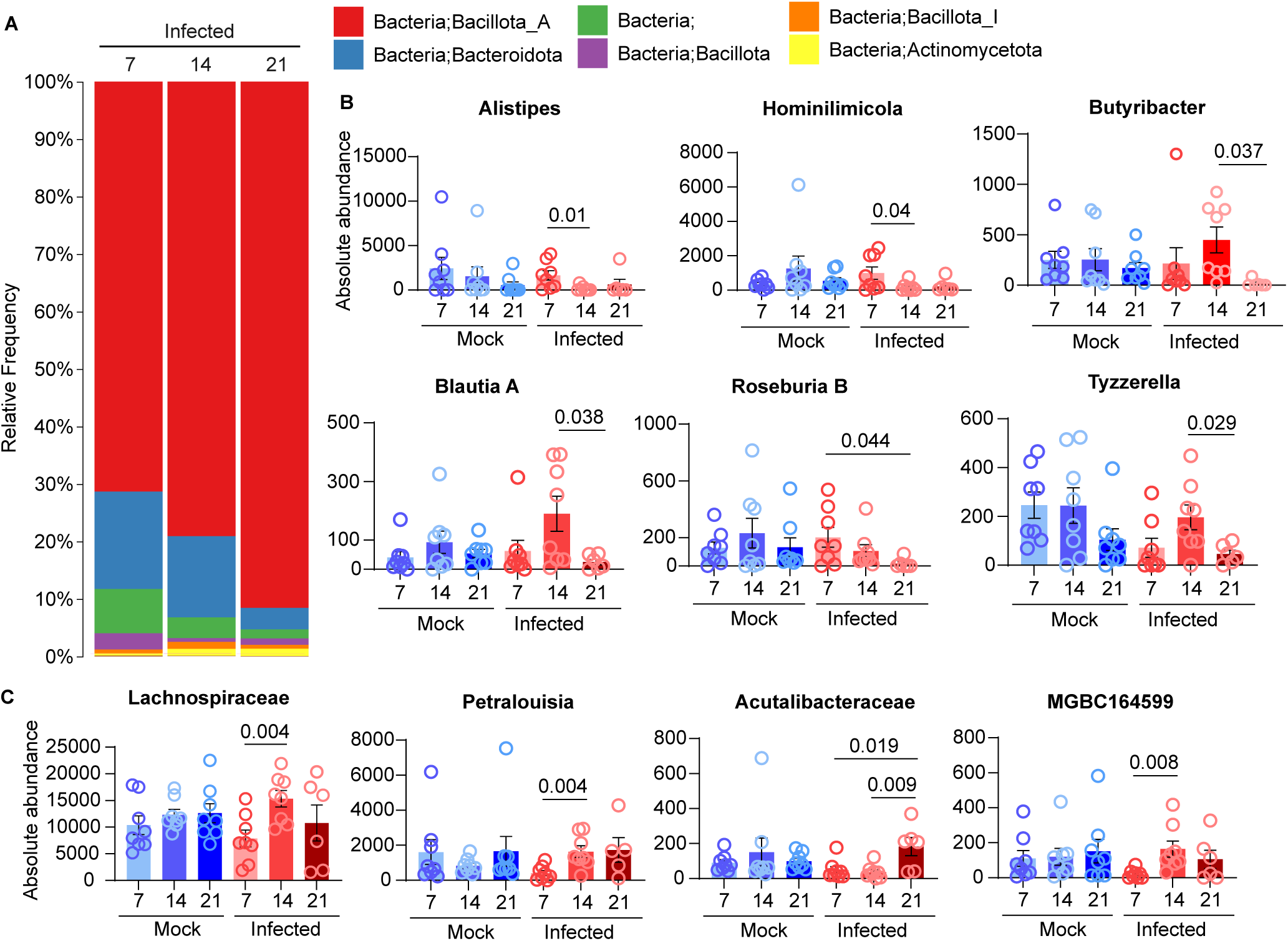
Changes over time in the post COVID phase. **A** Mean relative frequency of identified bacterial phyla across infected samples at 7, 14 and 21 dpi shown as percentage bar plots. **B, C** Absolute abundance level changes of Genus level bacteria between mock and infected and over time in infected condition which either decrease (B) or increase (C). Bar plots show individual values of biological replicates with error bars depicting SEM. Kruskal-Wallis (B, C) test performed for statistical analysis.

### 5.3 Post COVID-19 phase at 21 dpi shows significant microbiome dysbiosis

Despite recovery from acute infection, gut microbiome composition at 21 dpi showed clear divergence between infected and mock-infected mice. Alpha-diversity analyses revealed significant reductions in both Shannon diversity (p = 0.0127) and Pielou evenness (p = 0.0047) in infected animals, indicating a less diverse and less evenly distributed microbial community. In contrast, Chao1 richness did not differ significantly between groups (p = 0.3450), suggesting that total estimated species richness was largely preserved (**Fig. 3A**). To assess between-sample differences, we evaluated beta diversity using Bray–Curtis dissimilarity (**Fig. S1A**). Infected mice exhibited significantly altered community structure relative to mock controls at 14 dpi (p = 0.00375) and 21 dpi (p = 0.02) post-infection. Longitudinal comparisons within the infected group revealed no significant difference between 7 and 14 dpi, but pronounced divergence emerged by 21 dpi, with significant differences relative to both 7 dpi (p = 0.015) and 14 dpi (p = 0.0075). As expected, infected vs. mock at 7 dpi samples did not differ significantly, consistent with earlier findings (**Fig. S1A**). Within the mock group, longitudinal comparisons showed no significant differences across time points. Taxonomic profiling at 21 dpi demonstrated a continued expansion of Bacillota and loss of Bacteroidota in infected mice. Bacillota increased from 75.1% in mock-infected controls to 91.5% in infected animals, while Bacteroidota decreased from 19.4% to 3.7%, respectively (**Fig. 3B**). This pronounced phylum-level restructuring reflects a sustained reorganization toward a Bacillota-dominated community, consistent with the reduced evenness and altered beta-diversity patterns observed at this later time point. Notably, *Lawsonibacter* exhibited increased abundance in infected mice (**Fig. 3C**).

**Fig. 3.**
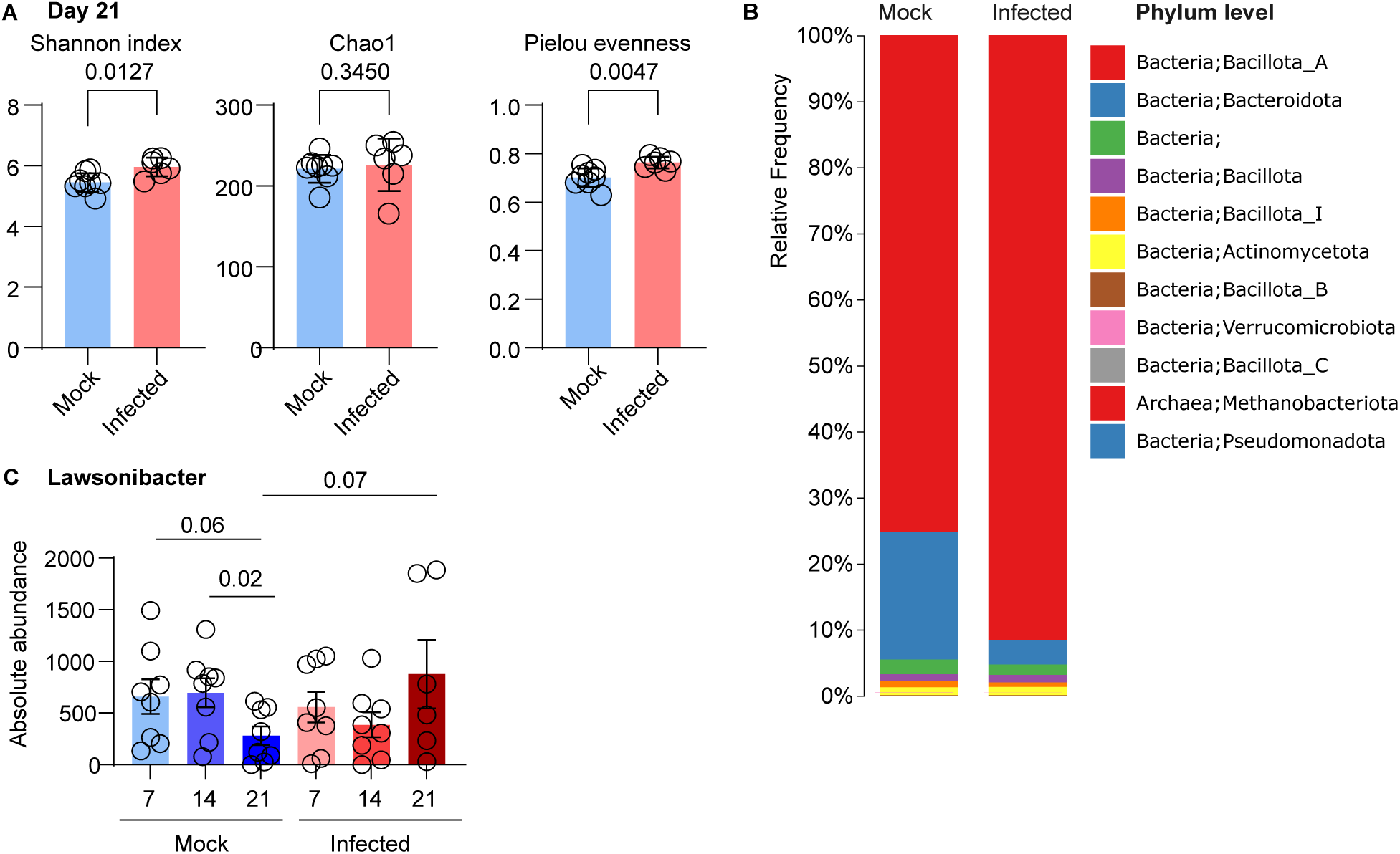
Microbiome changes at 21 dpi. **A** Shannon index, Chao1 and Pielou Evenness (Alpha Diversity) for mock and infected animals at 21 dpi. **B** Mean relative frequencies of identified bacterial phyla across mock and infected samples at 21 dpi shown as percentage bar plots. **C** Absolute abundance level changes of Genus level bacteria between mock and infected at 21 dpi. Bar plots show individual values of biological replicates with error bars depicting SEM. Mann-Whitney (A, D) and Kruskal-Wallis (B) test performed for statistical analysis.

### 5.4 Progressive Remodeling of Microbial Pathways

To determine whether SARS-CoV-2–associated shifts in microbial composition translated into functional changes, we performed PICRUSt pathway analysis at 7, 14, and 21 dpi. The number and magnitude of significantly altered pathways increased progressively over time from 7 dpi (15 pathways) to 14 dpi (17 pathways), and almost double at 21 dpi, set of 32 significantly altered pathways (**Fig. 4**). This escalation in functional disruption parallels the more pronounced taxonomic divergence observed at 21dpi. Using a combination of highest confidence level and smallest p-value the three most significantly altered pathways between mock and infected at 7 dpi are purine nucleotides degradation II (p=0.0038), mixed acid fermentation (p=0.038) and thiamin salvage II (p=0.01), indicating early changes in nucleotide turnover, fermentation capacity, and cofactor recycling. At 14 dpi, the functional profile shifted, with the most significantly affected pathways involving nitrate reduction VI (p=0.0012), thiamin salvage II (p=0.0045) and flavin biosynthesis I (0.0002). The recurrence of thiamin and flavin pathways at both 7 and 14 dpi indicates sustained alterations in microbial cofactor metabolism. At 21 dpi, the pathway landscape expanded markedly. The most significantly altered pathways included thiazole biosynthesis I (p=0.002), mixed acid fermentation (p=0.025) and the pathway of GDP-mannose-derived O-antigen building blocks biosynthesis (p=0.025). Numerous additional pathways such as the TCA cycle I (p=0.0093) and fucose and rhamnose degradation (p=0.0079) were also significantly different, indicating widespread remodeling of microbial metabolic potential at this late time point. Cross-timepoint comparison revealed clear patterns of overlap. Day 7 and 14 post-infection shared changes in 6-hydroxymethyl-dihydropterin diphosphate biosynthesis, flavin biosynthesis, and thiamin salvage, suggesting persistent disruption of folate-linked and redox-active cofactor pathways. Further, 7 and 21 dpi timepoints shared alterations in mixed acid fermentation and purine nucleotides degradation, indicating that fermentation-derived metabolites and purine turnover remain consistently affected across the followed timeline. Together, these findings demonstrate that SARS-CoV-2 infection drives progressively expanding functional alterations in the gut microbiome, with early disruptions in nucleotide and cofactor metabolism evolving into broad metabolic remodeling by 21 dpi.

**Fig. 4.**
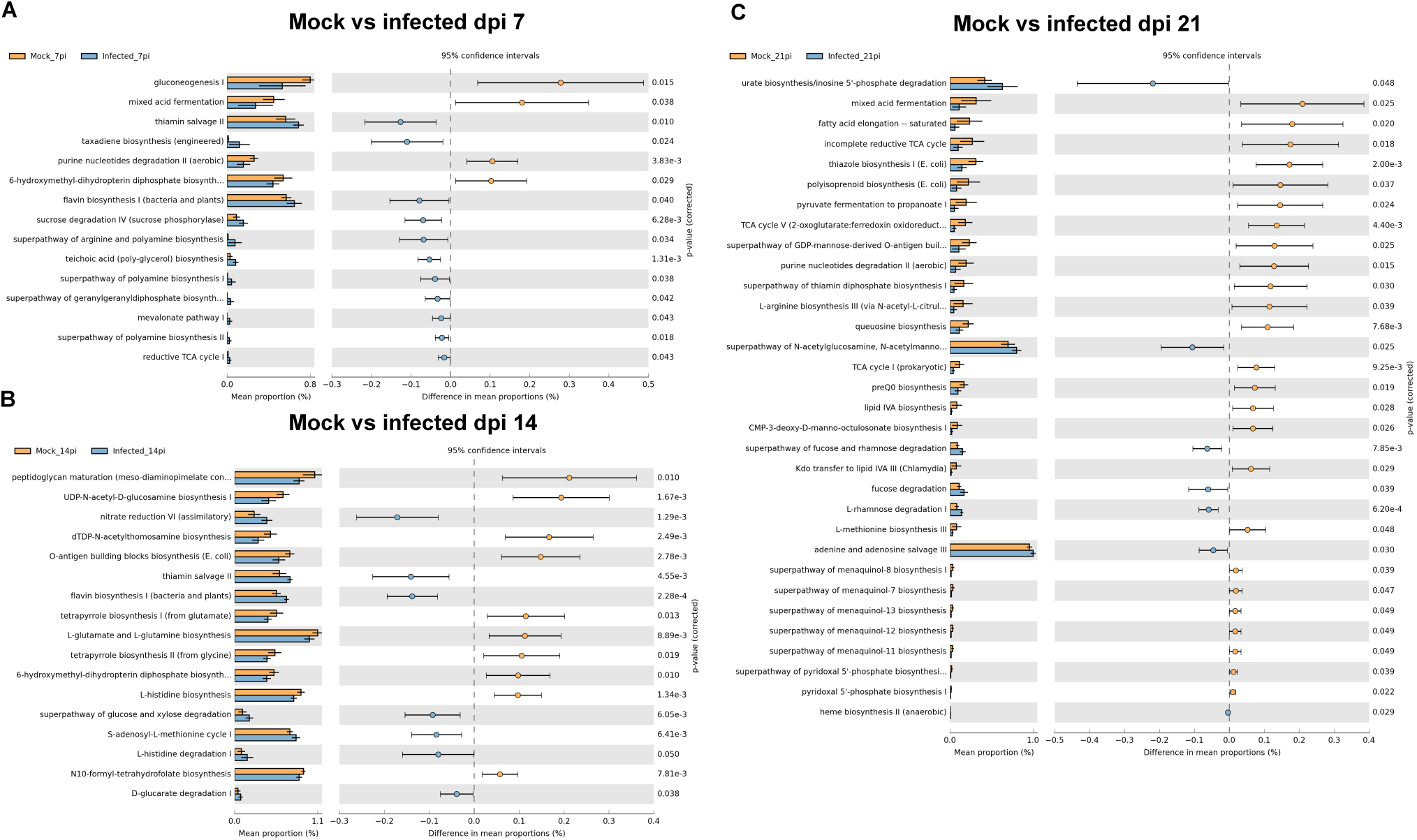
Picrust Pathway analyses at different timepoints. Error bar plots of identified pathways comparing mock vs infected mean proportion at 7 dpi (**A**), 14 (**B**), and 21 (**C**) of PICRUSt predicted MetaCyC function data, A Welch’s t-test for two groups was used for statistical analysis. An extended error bar plot was used to show differences in mean proportions of predicted functions with p < 0.05 (right y axis) between mock and corresponding infected group. The dop plot display differences using p-values. Values to the right of zero indicate pathways enriched in mock samples, while values to the left indicate enrichment in infected samples. Horizontal lines represent 95 % confidence intervals around the mean difference.

## 6. Discussion

This study demonstrates that SARS-CoV-2 infection produces a series of long-lasting alterations across the gut microbiome, extending beyond the period of acute viral replication in the lung. Microbial and functional pathway analyses across multiple time points revealed a trajectory of progressive dysbiosis over a period that coincides with the development of PCC symptoms.

Viral RNA dynamics confirmed a typical acute infection profile, with high pulmonary viral loads until 7 dpi, followed by a sharp decline and persistence of low-level RNA through 21 dpi. Alpha diversity, a measure of microbial richness and evenness, remained stable at 7 (end of acute phase) and 14 (early recovery) dpi but was significantly altered at 21 (PCC) dpi. This suggests that while infection does not drastically alter overall microbial richness, viral exposure contributes to later-stage microbiota alterations. In contrast, beta diversity, which reflects differences in microbial composition between infected and corresponding mock mice, showed substantial disruption at all time points, indicating that SARS-CoV-2 infection has a persistent impact on gut microbial community structure even after the transition from acute infection through recovery into PCC.

Despite the absence of early differences in alpha-diversity at 7 dpi, compositional remodeling was already evident at the phylum level, with an expansion of Bacillota and a reduction of Bacteroidota. These early changes increased over time, as longitudinal analyses revealed a two-phase pattern: an initial transition between 7 and 14 dpi involving taxa such as *Alistipes*, *Hominilimicola*, *Lachnospiraceae*, and *Petralouisia*, followed by a second wave of alterations emerging after 14 dpi. The latter phase was characterized by significant changes in additional Bacillota-associated genera, including *Butyribacter*, *Blautia A*, *Roseburia B*, *Tyzzerella*, and *Acutalibacteraceae*. By 21 dpi, the microbiome displayed pronounced dysbiosis, with reduced Shannon diversity and evenness, marked expansion of Bacillota, and near-complete loss of Bacteroidota. These findings indicate that SARS-CoV-2 infection initiates a progressive restructuring of the gut microbiome that becomes most pronounced at 21 dpi. These observations align with human studies reporting a shift toward Bacillota dominance in COVID-19 patients, a pattern proposed to contribute to systemic inflammation and immune dysregulation^18^. Conversely, the reduction in Bacteroidota, an important phylum for maintaining gut homeostasis and immune modulation^19^, may exacerbate prolonged inflammatory responses in PCC.

Recent work using the same mouse-adapted SARS-CoV-2 model has similarly identified delayed biological changes emerging around 21 dpi, including persistent viral RNA in multiple brain regions, elevated brainstem IL-6 expression, and a transition from depressive-to anxiety-like behaviors^16^. These findings demonstrate that post-acute neuroimmune disturbances continue to evolve well beyond the period of active viral replication. The convergence between late-phase brain alterations in that study and the pronounced gut microbial collapse observed here suggests that SARS-CoV-2 triggers a coordinated, multi-system shift becoming most pronounced at PCC stage. Such parallel trajectories raise the possibility that gut dysbiosis may contribute to, or be shaped by, the same inflammatory and neuroimmune processes implicated in PCC-related behavioral outcomes.

The microbial changes over time revealed patterns in post-infection dysbiosis. Notably, the bacterial taxa showing significant absolute abundance changes were primarily members of Bacillota and the order Clostridia, except for *Alistipes*, which belongs to Bacteroidota. This highlights the dominance of Clostridia-associated bacteria in shaping post-infectious microbiome restructuring. The observed fluctuations in bacterial populations further indicate a phased microbial adaptation, where certain taxa (e.g., *Acutalibacteraceae*, *Lawsonibacter*) expand later in infection, potentially compensating for earlier microbial losses, while others (e.g. *Alistipes*, *Hominiventricola*) are depleted early. These time-dependent bacterial shifts suggest that the gut microbiome undergoes a structured yet fluctuating response to SARS-CoV-2 infection rather than a linear return to homeostasis.

Functional pathway analysis showed that SARS-CoV-2 infection produced early disruptions in nucleotide turnover, fermentation, and cofactor metabolism, including thiamin and flavin pathways, which expanded into broader metabolic remodeling by 21 dpi. Across time points, pathways such as 6-hydroxymethyl-dihydropterin diphosphate biosynthesis, flavin biosynthesis, thiamin salvage, mixed acid fermentation, and purine nucleotide degradation were significantly altered, indicating sustained effects on microbial vitamin production and energy metabolism. By 21 dpi, additional changes in the TCA cycle and carbohydrate degradation reflected a shift toward widespread functional reprogramming of the microbiome. Disruption of vitamin-related pathways has been linked to metabolic deficiencies^20^, prolonged inflammation^21^, and cognitive impairment^22^, suggesting potential relevance for PCC-associated symptoms. Several pathways altered in this model including nitrogen cycling, amino acid metabolism, purine nucleobase degradation, and histidine degradation mirror findings in human COVID-19 cohorts^23^, supporting the translational relevance of the functional trajectory observed here, using this murine model.

Several limitations should be considered when interpreting these findings. First, microbiome analyses relied on 16S rRNA sequencing, which limits taxonomic resolution and does not directly assess microbial metabolites or host–microbe interactions. Second, peak viral replication in our model occurs at 2–4 dpi, with 7 dpi marking the end of the acute phase, 14 dpi reflecting recovery, and 21 dpi corresponding to PCC. Because pre-infection baselines and early acute-phase samples were not included, future studies will be required to fully resolve the directionality, magnitude, and timing of microbiome disruptions. Finally, the study focused on a 21-day window, and longer-term follow-up will be necessary to establish whether microbiome and behavioral alterations persist, resolve, or evolve beyond this period.

Taken together, these findings outline a clear post-infection trajectory in which SARS-CoV-2 drives progressive gut microbiome dysbiosis, expanding functional disruptions, and sensory changes persisting up to 21 dpi, at least. Identifying these microbial signatures may provide novel biomarkers for predicting PCC development risk and severity, as well as offer potential targets for microbiome-based interventions aimed at restoring gut homeostasis. Microbiome-targeted interventions, such as prebiotics and probiotics, may thus aid in recovery from COVID-19, mitigating the risk of PCC and improve immune resilience.

## 7. Materials and Methods

### 7.1 Animal model

All animal procedures were approved by the University of Alberta Animal Care and Use Committee (AUP00003963) and conducted in accordance with institutional guidelines. Twenty-three-to twenty-five-week-old male C57BL/6J mice (Jackson Laboratory, Bar Harbor, ME, USA; strain #000664) were housed in a BSL-3 facility in groups of up to five per ventilated cage with ad libitum access to food (5L0D, LabDiet, St. Louis, MO, USA) and water. Animals were maintained on a 12-h light–dark cycle at 22 °C and 52% humidity, and health and body weight were monitored daily. Mice were infected intranasally under isoflurane anesthesia (Fresenius Kabi, Bad Homburg, Germany) with 1 × 10⁴ plaque-forming units (PFU) of mouse-adapted SARS-CoV-2/3AA^24^ in 50 µL Dulbecco’s Modified Eagle Medium (DMEM; Gibco). Control animals received 50 µL serum-free DMEM. At 2, 7, 14, and 21 days post-infection (dpi), animals were euthanized by isoflurane overdose followed by cardiac puncture. Tissue collection adhered to the methodology established by Xu et al.^25^. In brief, the left lung lobe was excised and homogenized in 1 mL DMEM using a GentleMACS™ Dissociator (Miltenyi Biotec, Bergisch Gladbach, Germany) and GentleMACS™ M Tubes. Homogenates were centrifuged for 10 min at 4°C at 4,000 × g. Cleared supernatants were aliquoted and stored at −80 °C. Viral RNA in lung was quantified by droplet digital PCR as described in Roczkowsky et al.^26^.

### 7.2 Feces Collection and processing

During necropsy, the small intestine (from the pylorus to the caecum) was excised and placed in ice-cold sterile phosphate-buffered saline (PBS; Thermo Fisher Scientific, Waltham, MA, USA) on a sterile Petri dish. The duodenum (proximal quarter), jejunum (middle half), and ileum (distal quarter) were separated. Intestinal contents were flushed using a 30 mL syringe (BD, Franklin Lakes, NJ, USA) filled with ice-cold PBS and by gently expressing luminal material with a 23-gauge blunted needle (BD, Franklin Lakes, NJ, USA). Stool was collected into 1.5 mL microcentrifuge tubes (Eppendorf, Hamburg, Germany), snap-frozen in liquid nitrogen, and stored at −80 °C until processing. For DNA extraction, frozen stool was transferred into 2 mL tubes containing 2.8 mm ceramic beads (Thermo Fisher Scientific, Waltham, MA, USA) and a 1:1 (v/v) mixture of freshly prepared lysis buffer (proteinase K, RNase A, and Buffer ATL from the DNeasy® Blood & Tissue Kit; Qiagen, Hilden, Germany; Cat. 69506) and nuclease-free water (Thermo Fisher Scientific, Waltham, MA, USA). Samples were homogenized by vortexing for 1 min and heat-inactivated at 70 °C for 30 min using a Thermo Fisher Scientific dry-block heater. Genomic DNA was isolated using the DNeasy® Blood & Tissue Kit according to the manufacturer’s protocol. DNA concentration and purity were assessed using a NanoDrop™ ND-1000 spectrophotometer (Thermo Fisher Scientific, Waltham, MA, USA).

### 7.3 16S DNA sequencing of microbiome

Samples were submitted to Novogene Corporation Inc. (Sacramento, CA, USA) for 16S rRNA gene amplicon sequencing targeting the V3–V4 region, generating approximately 30,000 paired-end reads per sample. Raw FASTQ files were processed in-house. Primer trimming was performed using Cutadapt and read quality was assessed using FastQC^27^. Reads were filtered at phred ≥ 30. Amplicon sequence variants (ASV) were inferred using DADA2^28^ with error-model learning, dereplication, paired-end merging, and chimera removal. Taxonomic assignment was performed using the Genome Taxonomy Database (GTDB) reference database, release 09-RS220. ASV tables were rarefied to a depth of 47,055 reads prior to diversity analyses. Beta-diversity metrics (Bray–Curtis), PERMANOVA tests, visualization of taxonomic composition and alpha diversity metrics were computed in QIIME2^29^ 2023.9. Functional pathway predictions were generated using PICRUSt2^30^. All sequencing data have been deposited in the NCBI BioProject database under accession number PRJNA1236398.

### 7.4 Statistical analysis

All statistical analyses for visualizations were performed in GraphPad Prism version 10.0.1 (GraphPad Software, San Diego, CA, USA). Data distributions were assessed for normality using the Shapiro–Wilk test, which revealed that all datasets were non-normally distributed. Accordingly, Mann–Whitney U tests for two-group comparisons and Kruskal–Wallis tests for comparisons involving more than two groups were used. The specific statistical test used for each dataset is indicated in the corresponding figure legend. All bar graphs display mean ± SEM.

## 9. Supplementary figures and tables

**Supplementary Figures. Fig. S1. Beta Diversity Analysis.** Bray-Curtis Distance shown as NMDS (Non-metric multi-dimensional scaling) plot (left) using all time points of infected and mock samples, with the table showing corresponding statistical parameters (right).

**Supplementary Tables. Table S1:** Absolute abundance of identified bacterial phyla, class, order, family genus and species.

## 10. Author statements

### 10.1 Author Contributions

Conceptualization, C.P., D.O., M.A., M.E., N.O., P.F., R.M., S.M.; methodology, M.A., N.O., R.M.; software, R.M., S.M.; formal analysis, N.O., R.M., S.M.; investigation, D.O., M.E., N.O., R.M.; resources, C.P., P.F.; data curation, R.M.; writing—original draft preparation, R.M.; writing—review and editing, N.O., P.F.; visualization, N.O., R.M., S.M.; supervision, C.P., P.F.; project administration, N.O., P.F., R.M.; funding acquisition, C.P., P.F. All authors have read and agreed to the published version of the manuscript.

### 10.4 Acknowledgments

P.F. is the AstraZeneca (Canada) Chair in Asthma and Obstructive Lung Disease. C.P. is supported by the H.M. Toupin Chair in Neurocognitive Disorders.

