## Supplementary material for "Progressive Post-Acute Gut Microbiome Disruption Following SARS-CoV-2 Infection in Mice": Fig. S1

A

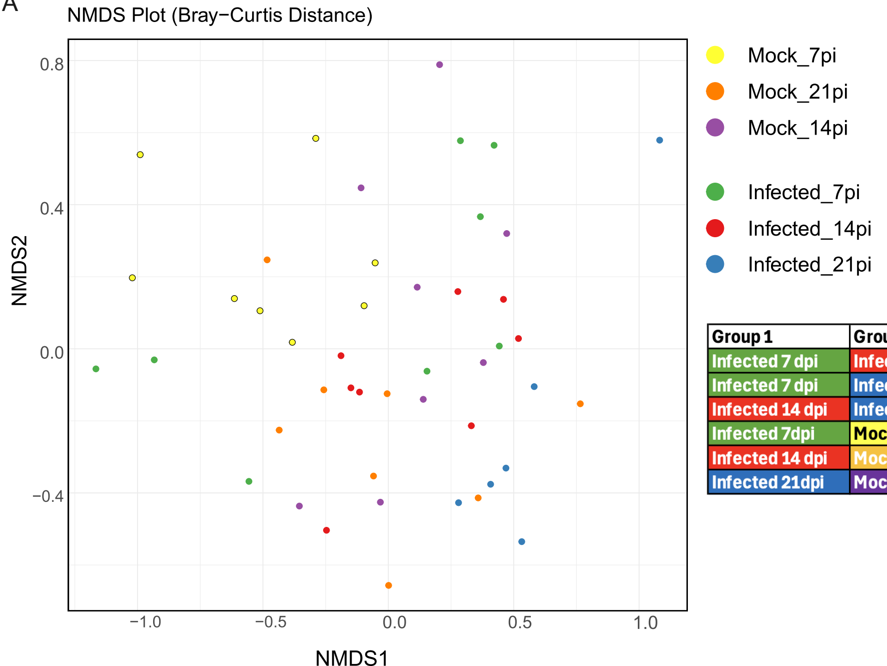


| Group 1 | Group 2 | Sample size | Permutations | pseudo-F | p-value |
|---|---|---|---|---|---|
| Infected 7 dpi | Infected 14 dpi | 16 | 999 | 2.34 | 0.036 |
| Infected 7 dpi | Infected 21 dpi | 14 | 999 | 2.89 | 0.007 |
| Infected 14 dpi | Infected 21 dpi | 14 | 999 | 2.76 | 0.003 |
| Infected 7dpi | Mock 7 dpi | 16 | 999 | 1.87 | 0.068 |
| Infected 14 dpi | Mock 14 dpi | 16 | 999 | 2.51 | 0.001 |
| Infected 21dpi | Mock 21 dpi | 14 | 999 | 2.13 | 0.012 |
